# The ARRIVE guidelines 2019: updated guidelines for reporting animal research

**DOI:** 10.1101/703181

**Authors:** Nathalie Percie du Sert, Viki Hurst, Amrita Ahluwalia, Sabina Alam, Marc T. Avey, Monya Baker, William J. Browne, Alejandra Clark, Innes C. Cuthill, Ulrich Dirnagl, Michael Emerson, Paul Garner, Stephen T. Holgate, David W. Howells, Natasha A. Karp, Katie Lidster, Catriona J. MacCallum, Malcolm Macleod, Ole Petersen, Frances Rawle, Penny Reynolds, Kieron Rooney, Emily S. Sena, Shai D. Silberberg, Thomas Steckler, Hanno Würbel

## Abstract

Reproducible science requires transparent reporting. The ARRIVE guidelines were originally developed in 2010 to improve the reporting of animal research. They consist of a checklist of information to include in publications describing *in vivo* experiments to enable others to scrutinise the work adequately, evaluate its methodological rigour, and reproduce the methods and results. Despite considerable levels of endorsement by funders and journals over the years, adherence to the guidelines has been inconsistent, and the anticipated improvements in the quality of reporting in animal research publications have not been achieved.

Here we introduce ARRIVE 2019. The guidelines have been updated and information reorganised to facilitate their use in practice. We used a Delphi exercise to prioritise the items and split the guidelines into two sets, the ARRIVE Essential 10, which constitute the minimum requirement, and the Recommended Set, which describes the research context. This division facilitates improved reporting of animal research by supporting a stepwise approach to implementation. This helps journal editors and reviewers to verify that the most important items are being reported in manuscripts. We have also developed the accompanying Explanation and Elaboration document that serves 1) to explain the rationale behind each item in the guidelines, 2) to clarify key concepts and 3) to provide illustrative examples. We aim through these changes to help ensure that researchers, reviewers and journal editors are better equipped to improve the rigour and transparency of the scientific process and thus reproducibility.

## Why good reporting is important

In recent years issues about the reproducibility of research findings have raised considerable concern among scientists, funders, research users and policy makers [1–3]. Important contributing factors identified include flawed study design and analysis, variability and inadequate validation of reagents and other biological materials, insufficient reporting of methodology and results, and barriers to access data [4]. A number of initiatives have been developed to improve the reproducibility of scientific research, from funders’ open access policies [5], through to alternative peer review models [6], and the development of infrastructure to promote study preregistration and data sharing [7].

Transparent reporting is an essential first step for any initiative focusing on reproducibility. Without this, the methodological rigour of the studies cannot be adequately scrutinised, the reliability of the findings cannot be assessed, and the work cannot be repeated or built upon by others. Despite the development of specific reporting guidelines for preclinical and clinical research, evidence suggests that scientific publications often lack key information and that there continues to be considerable scope for improvement [8–14]. Animal research is a good case in point, where poor reporting impacts on the development of therapeutics and irreproducible publications can spawn an entire field of research, or trigger clinical studies, subjecting patients to interventions unlikely to be effective [2, 15, 16].

In an attempt to improve the reporting of animal research, the ARRIVE guidelines were published in 2010. The guidelines consist of a checklist of the information that should be included in any manuscript describing animal-based research, to ensure that the research is described in a comprehensive and transparent manner [17–27]. In the nine years since publication, the ARRIVE guidelines have been endorsed by more than a thousand journals from across the life sciences. Endorsement typically includes advocating their use in guidance to authors and reviewers. However, only a small number of journals actively enforce compliance; recent studies have shown that important information as set out in the ARRIVE guidelines is still missing from most publications sampled. This includes randomisation (reported in only 30-40% of publications), blinding (reported in only approximately 20% of publications), sample size justification (reported in less than 10% of publications) and animal characteristics (all basic characteristics reported in less than 10% of publications) [28–30].

Evidence suggests two main factors limit the impact of the guidelines. The first is the extent to which editorial and journal staff are involved in enforcing reporting standards. A randomised controlled trial at PLOS ONE, for example, demonstrated that a request by journal staff to include a completed ARRIVE checklist in the manuscript submission process did not improve the disclosure of information in published papers [31]. In contrast, other studies using reporting checklists with more editorial follow up have shown a marked improvement in the nature and detail of the information included in publications [32–34]. Providing the level of journal or editorial input required to ensure compliance with all the items of the ARRIVE guidelines is unlikely to be sustainable for most journals because of the resources needed. Requesting adherence with all items at once, with no consideration of their relative importance might also be perceived as too prescriptive and further complicate the task.

The second issue is that researchers and other individuals and organisations responsible for the integrity of the research process are not sufficiently aware of the consequences of incomplete reporting. There is some evidence that awareness of ARRIVE is linked to the use of more rigorous experimental design standards [35], but there is also evidence that researchers are unaware of the much larger systemic bias in the publication of research and in the reliability of certain findings and even of entire fields [31, 36–38]. This lack of understanding affects how experiments are designed and grant proposals prepared, how animals are handled and data recorded in the laboratory, and how manuscripts are written by authors or assessed by journal staff, editors and reviewers.

Approval for experiments involving animals is generally based on a harm-benefit analysis, weighing the harms to the animals involved against the benefits of the research to society. If the research is not reported in enough detail, even when conducted rigorously, the benefits may not be realised, and the harm-benefit analysis and public trust in the research are undermined [39]. As a community, we must do better to ensure that where animals are used the research is well designed and analysed, and transparently reported. Here we introduce the revised ARRIVE guidelines, referred to as ARRIVE 2019. The information included has been updated, extended and reorganised to facilitate the use of the guidelines, helping to ensure that researchers, editors and reviewers, as well as other relevant journal staff, are better equipped to improve the rigour and reproducibility of animal research.

## Introducing ARRIVE 2019

The revision of the ARRIVE guidelines has been undertaken by a new international working group – the authors of this publication. Our expertise comes from across the life sciences community, including funders, journal editors, statisticians, methodologists and researchers from academia and industry. We have improved the clarity of the guidelines, prioritised the items, added new information and generated the accompanying Explanation and Elaboration document to provide context and rationale for each item [40]. New additions comprise inclusion and exclusion criteria, which are a key aspect of data handling and prevent the ad hoc exclusion of data [41]; protocol registration, a recently emerged approach which promotes scientific rigour and encourages researchers to carefully consider the experimental design and analysis plan before any data are collected [42]; and data access, in line with the FAIR Data Principles [43]. Table S1 summarises the changes.

The most significant departure from the original guidelines is the classification of items into two prioritised groups, as shown in Tables 1 and 2. There is no ranking within each group. The first group is the “ARRIVE Essential 10” which describes information that is the basic minimum to include in a manuscript, as without this information reviewers and readers cannot confidently assess the reliability of the findings presented. It includes the study design, sample size, measures to reduce subjective bias such as randomisation and blinding, outcome measures, statistical methods, experimental animals used, experimental procedures and results. The second group, referred to as the “Recommended Set” adds context to the study described. This includes the ethical statement, declaration of interest, protocol registration and data access, as well as more detailed information on the methodology such as animal housing, husbandry, care and monitoring. Items on abstract, background, objectives, interpretation and generalisability also describe what to include in the more narrative parts of a manuscript. The prioritisation was derived from a Delphi exercise [44] to rank items according to their relative importance for assessing the reliability of research findings. The Delphi panel included the working group in addition to external stakeholders, to ensure maximum diversity in fields of expertise and geographical location. Demographics of the Delphi panel and full methods and results are presented in Supporting Information S2 and S3.

**Table 1.** ARRIVE Essential 10

|  |  |  |
| --- | --- | --- |
| Study design | 1 | For each experiment, provide brief details of study design including:<br>a. The groups being compared, including control groups. If no control group has been used, the rationale should be stated.<br>b. The experimental unit (e.g. a single animal, litter, or cage of animals). |
| Sample size | 2 | a. Specify the exact number of experimental units allocated to each group, and the total number in each experiment. Also indicate the total number of animals used.<br>b. Explain how the sample size was decided. Provide details of any *a priori* sample size calculation, if done. |
| Inclusion and exclusion criteria | 3 | a. Describe any criteria established *a priori* for including and excluding animals (or experimental units) during the experiment, and data points during the analysis.<br>b. For each experimental group, report any animals, experimental units or data points not included in the analysis and explain why.<br>c. For each analysis, report the exact value of N in each experimental group. |
| Randomisation | 4 | Describe the methods used:<br>a. To allocate experimental units to control and treatment groups. If randomisation was used, provide the method of randomisation.<br>b. To minimise potential confounding factors such as the order of treatments and measurements, or animal/cage location. |
| Blinding | 5 | Describe who was aware of the group allocation at the different stages of the experiment (during the allocation, the conduct of the experiment, the outcome assessment, and the data analysis). |
| Outcome measures | 6 | a. Clearly define all outcome measures assessed (e.g. cell death, molecular markers, or behavioural changes).<br>b. For hypothesis-testing studies, specify the primary outcome measure, i.e. the outcome measure that was used to determine the sample size. |
| Statistical methods | 7 | a. Provide details of the statistical methods used for each analysis.<br>b. Specify the experimental unit that was used for each statistical test.<br>c. Describe any methods used to assess whether the data met the assumptions of the statistical approach. |
| Experimental animals | 8 | a. Provide details of the animals used, including species, strain and substrain, sex, age or developmental stage, and weight.<br>b. Provide further relevant information on the provenance of animals, health/immune status, genetic modification status, genotype, and any previous procedures. |
| Experimental procedures | 9 | For each experimental group, including controls, describe the procedures in enough detail to allow others to replicate them, including:<br>a. What was done, how it was done and what was used.<br>b. When and how often.<br>c. Where (including detail of any acclimation periods).<br>d. Why (provide rationale for procedures). |
| Results | 10 | For each experiment conducted, including independent replications, report:<br>a. Summary/descriptive statistics for each experimental group, with a measure of variability where applicable.<br>b. If applicable, the effect size with a confidence interval. |

**Table 2.** ARRIVE Recommended Set

| Recommended Set |  |  |
| --- | --- | --- |
| Abstract | 11 | Provide an accurate summary of the research objectives, animal species, strain and sex, key methods, principal findings, and study conclusions. |
| Background | 12 | a. Include sufficient scientific background to understand the rationale and context for the study, and explain the experimental approach.<br>b. Explain how the animal species and model used address the scientific objectives and, where appropriate, the relevance to human biology. |
| Objectives | 13 | Clearly describe the research question, research objectives and, where appropriate, specific hypotheses being tested. |
| Ethical statement | 14 | Provide the name of the ethical review committee or equivalent that has approved the use of animals in this study and any relevant licence or protocol numbers (if applicable). If ethical approval was not sought or granted, provide a justification. |
| Housing and husbandry | 15 | Provide details of housing and husbandry conditions, including any environmental enrichment. |
| Animal care and monitoring | 16 | a. Describe any interventions or steps taken in the experimental protocols to reduce pain, suffering and distress.<br>b. Report any expected or unexpected adverse events.<br>c. Describe the humane endpoints established for the study and the frequency of monitoring. |
| Interpretation /scientific implications | 17 | a. Interpret the results, taking into account the study objectives and hypotheses, current theory and other relevant studies in the literature.<br>b. Comment on the study limitations including potential sources of bias, limitations of the animal model, and imprecision associated with the results. |
| Generalisability /translation | 18 | Comment on whether, and how, the findings of this study are likely to generalise to other species or experimental conditions, including any relevance to human biology (where appropriate). |
| Protocol registration | 19 | Provide a statement indicating whether a protocol (including the research question, key design features, and analysis plan) was prepared before the study, and if and where this protocol was registered. |
| Data access | 20 | Provide a statement describing if and where study data are available. |
| Declaration of interests | 21 | a. Declare any potential conflicts of interest, including financial and non-financial. If none exist, this should be stated.<br>b. List all funding sources (including grant identifier) and the role of the funder(s) in the design, analysis and reporting of the study. |

The classification of the items into two groups is intended to facilitate the improved reporting of animal research by allowing an initial focus on the most critical issues. This better allows journal staff, editors and reviewers to verify that the items have been adequately reported in manuscripts. The first step should be to ensure compliance with the ARRIVE Essential 10 as a minimum requirement. Items from the Recommended Set can then be added over time and in line with specific editorial policies until all the items are routinely reported in all manuscripts.

Although the guidelines are written with researchers and journal editorial policies in mind, it is important to stress that researchers alone should not have to carry the burden of responsibility for transparent reporting. Funders, institutions and publishers all have a responsibility to ensure that the appropriate training, workflows and practices are in place to support researchers in their different roles. In particular, institutions and other research performing organisations, both public and private, as well as publishers, have a key role to play in building capacity and championing the behavioural changes required to improve reporting practices.

## Conclusion

Transparent reporting is clearly essential if animal studies are to add to the knowledge base and inform future research, policy and clinical practice. ARRIVE 2019 prioritises the reporting of information related to study reliability. This enables research users to assess how much weight to ascribe to the findings, and in parallel promotes the use of rigorous methodology in the planning and conduct of *in vivo* experiments [35], thus increasing the likelihood that the findings are reliable, and ultimately, reproducible.

The intention of ARRIVE 2019 is not to supersede individual journal requirements but to promote a harmonised approach across journals to ensure that all manuscripts contain essential information needed to appraise the research. The step-by-step approach is in line with current practice by a number of journals who recommend that authors not only refer to the ARRIVE guidelines while preparing their manuscript, but also implement a checklist with a core set of items. Journals usually share a common objective of improving the methodological rigour and reproducibility of the research they publish, but different journals emphasise different pieces of information [45–47]. Here we propose an expert consensus on information to prioritise. This will provide clarity for authors, facilitate transfer of manuscripts between journals, and accelerate an improvement of reporting standards.

Concentrating the efforts of the research and publishing communities on the ARRIVE Essential 10 items provides a manageable approach to evaluate reporting quality efficiently and assess the effect of interventions and policies designed to improve the reporting of animal experiments. It also provides a starting point for the development of automated or semi-automated artificial intelligence tools that can detect missing information rapidly [48].

Improving reporting is a collaborative endeavour and concerted effort from the biomedical research community is required to ensure maximum impact. We welcome collaboration with other groups operating in this area, and feedback on ARRIVE 2019 and our implementation strategy.

## Supporting information

S1 Table: Noteworthy changes in ARRIVE 2019, compared to ARRIVE 2010

S2 Delphi methods and results

S3 Delphi data

## Acknowledgements

We would like to thank the members of the expert panel for the Delphi exercise, and the DelphiManager team for advice and use of their software. We would like to acknowledge the late Doug Altman’s contribution to this project, Doug was a dedicated member of the working group and his input to the guidelines’ revision has been invaluable.

---

**S1 Table:** Noteworthy changes in ARRIVE 2019, compared to ARRIVE 2010

| ARRIVE 2019 | ARRIVE 2010 | Reason for change |
| --- | --- | --- |
| All items | All items | We reordered items and split them in two sets based on their importance to assess the reliability of the study. There is no ranking within each set, items are ordered logically. |
| **ARRIVE Essential 10** |  |  |
| Item 1 – Study design | Item 6 – Study design | We removed the reference to steps taken to minimise the effects of bias (formerly subitem 6b). All information about randomisation is now in item 4 and all information about blinding is now in item 5. |
| Item 2 – Sample size | Item 10 – Sample size | We clarified that the number of experimental units might be different from the number of animals. Independent replications are now mentioned with the results (item 10) to prevent any confusion with biological replicates. |
| Item 3 – Inclusion and exclusion criteria | Item 15 – Numbers analysed | We added a new subitem on a priori inclusion and exclusion criteria, evidence shows that ad hoc exclusion of data can lead to false positive results [1]. We clarified that the N number in each analysis might be different from the number of animals. We renamed the item to better reflect content. |
| Item 4 – Randomisation | Item 11 – Allocating animals to experimental groups | All references to randomisation were consolidated in this item for clarity. We reworded the text to include the randomisation procedure which was covered separately in the study design (formerly item 6). We clarified that experimental units are allocated to group, rather than animals. |
| Item 5 – Blinding | Item 6 – Study design | Blinding was included in the original guidelines as part of the study design (formerly subitem 6b), we have added more text in a new item to highlight its importance and encourage greater specificity. |
| Item 6 – Outcome measures | Item 12 – Experimental outcomes | We clarified that all outcome measures should be reported, and added a subitem to highlight the need to identify a primary outcome measure for hypothesis-testing studies.<br>We changed the item name to 'outcome measures' because of concerns within the group that the term 'experimental outcomes' could be ambiguous. |
| Item 7 – Statistical methods | Item 13 – Statistical methods | We removed a reference to unit of analysis, which is often poorly understood and used the term experimental unit for consistency throughout the guidelines. |
| Item 8 – experimental animals | Item 8 – experimental animals | We clarified the wording and removed examples to streamline the guidelines, further details are discussed in the supporting E&E document [2]. |
| Item 9 – Experimental procedures | Item 7 – Experimental procedures | We encouraged greater specificity by stating that procedures should be described in enough detail to allow others to replicate them. We removed examples to streamline the guidelines, further details are discussed in the supporting E&E document [2]. |
| Item 10 – Results | Item 16 – outcomes and estimation | We expanded this item to provide more explicit guidance on reporting results. The name of the item was changed from 'outcomes and estimations' to 'results' for clarity and prevent confusion with item 6 – outcome measures. |
| Item removed | Item 14 – Baseline data | This item overlapped with item 8 – Experimental animals and the two items were combined, with further details provided in the supporting E&E document [2]. |

| ARRIVE Recommended Set |  |  |
| --- | --- | --- |
| Item 11 – Abstract | Item 2 – Abstract | We specified that the sex of animals used should be included the abstract, empirical evidence suggests an endemic male bias in biomedical research [3]. |
| Item 12 – Background | Item 3 – Background | We clarified the wording and removed examples to streamline the guidelines, further details are discussed in the supporting E&E document [2]. |
| Item 13 – Objectives | Item 4 – Objectives | We removed a reference to primary and secondary objectives as it would not apply to exploratory studies, and added a requirement to describe the research question, which is relevant to all study types. |
| Item 14 – Ethical statement | Item 5 – Ethical statement | We removed reference to UK legislation to make this item relevant for an international audience. We added specification of the relevant licence or protocol numbers to provide accountability and promote transparency. |
| Item 15 – Housing and husbandry | Item 9 – Housing and husbandry | We moved the subitem on welfare-related assessments and interventions to item 16 – Animal care and monitoring. We removed examples to streamline the guidelines, further details are discussed in the supporting E&E document [2]. |
| Item 16 – Animal care and monitoring | Item 17 – Adverse events | We added a new subitem to encourage the reporting of humane endpoints and monitoring, and changed the name of the item to animal care and monitoring to better reflect content. |
| Item 17 – Interpretation/scientific implications | Item 18 – Interpretation/scientific implications | We removed subitem c “Describe any implications of your experimental methods or findings for the replacement, refinement or reduction (3Rs) of the use of animals in research”. This item is not relevant to all animal studies and further details have been provided in the supporting E&E document [2]. |
| Item 18 – Generalisability/translation | Item 19 – Generalisability/translation | We simplified and clarified the wording. |
| Item 19 – Protocol registration | New item | We added a new item on registering key aspects of the protocol. Empirical studies have shown up to 50% of outcomes which are measured are not reported [4]. This selective outcome reporting bias leads to an overstatement of biological effects. |
| Item 20 – Data access | New item | We added a new item on data access to encourage authors to provide a data sharing statement describing how others can gain access to the data on which the paper is based. |
| Item 21 – Declaration of interests | Item 20 – Funding | We added a new sub-item on declaring potential conflicts of interest. We added the specification of the role of the funder(s) in the ‘design, analysis and reporting of the study’. This information allows the reader to assess any competing interests, and any potential sources of bias.<br>We renamed the item to better reflect content. |
| Item removed | Item 1 – Title | We removed this item as it provided no specific guidance on what to include in the title. |

## S2 Delphi methods and results

### 1 Methods

#### 1.1 Development of the revised checklist

For each subitem of ARRIVE 2010, the NC3Rs summarised the evidence justifying its inclusion in the guidelines and any indication of a need for revision. The working group then met for a two-day meeting in November 2017 in London, to review this information, discuss the addition of new items and agree on the strategy to go forward. We agreed to update the guidelines, develop an explanation and elaboration (E&E) document for ARRIVE 2019 [1] and prioritise the items to facilitate the uptake of the revised guidelines [2]. After the meeting, each item was allocated to at least two members of the working group to develop the item’s explanation in more detail and refine the item’s wording. Further iterations of the checklist were achieved by email discussion within the whole group.

The Delphi exercise was designed to achieve consensus on prioritising items of the ARRIVE guidelines.

The objective was to allocate the 22 items into two or three shortlists with different levels of priority, and relatively even distribution within each set.

#### 1.2 Recruitment of the Delphi expert panel

Ethical approval for this study was obtained from the University of Bristol, Faculty of Science Research Ethics Committee (ID 66625).

The panel consisted of the ARRIVE Working Group and external stakeholders nominated by the Working Group, with suitable expertise on the quality of animal research or its reporting. We aimed to gather a diverse panel of experts, both in terms of field of expertise and geographical location.

Panel members consented to take part by following a link in the invitation email to the first round.

#### 1.3 The Delphi process

There were three iterations of the questionnaire in total [3], and these were managed using the Comet Initiative DelphiManager platform (http://www.comet-initiative.org/DelphiManager/). Data collection took place June to November 2018. Panel members received an email invitation at the start of each round with a link to the online questionnaire. They were allowed three weeks to complete the questionnaire, with email reminders at day 7 and day 14. If they did not respond within the time frame they were excluded from that round, however they were invited to take part in the subsequent rounds of the Delphi.

Each of the 22 items of the revised ARRIVE guidelines was evaluated against the statement:

**“How important is this piece of information for assessing the reliability of results in an animal research paper?”**

Panel members scored each item on a scale of 1 – 9, where 1 was least important and 9 was most important.

The questionnaire presented in round 1 included free-text fields to provide reasoning for the score given to each item. Individual justifications were collated, summarised and presented to the whole panel in round 2.

In round 2, panel members were asked to provide a justification if their score for a particular item had changed between round 1 and round 2. Similarly, this information was summarised and presented to the whole panel in round 3.

Following rounds 1 and 2, the scores for each item were analysed and a structured summary consisting of a histogram showing the dispersal of the scores in the entire panel was prepared. This summary was presented with a new iteration of the questionnaire at the next round, where panel members were asked to re-score the items. In round 2 and 3, panel members’ own scores from the previous round were also displayed for each item.

To encourage a wider dispersal of scores, in the final round (round 3) panel members were asked to follow two rules while scoring items:

- to score no more than ten items in the top range (7 – 9)
- to score no fewer than six items in the bottom range (1 – 3)

In the final dataset we excluded data entries which had not followed these rules, allowing for a deviation of ±1 item in each range.

#### 1.4 Addition of new ARRIVE items

In the first round of the Delphi, we asked the panel to suggest new items that they believed should be included in the revised guidelines. The threshold for inclusion was defined a priori; for a new item to be considered, it would have to be suggested by at least 10% of the panel. The panel also had the opportunity to provide general feedback at the end of the survey.

#### 1.5 Criteria for allocating items to sets

The plan to achieve consensus was defined a priori and two options were considered. The first option was to allocate the items in three sets, based on each item’s median score and a minimum of 70% of the panel scoring the item within the same range. Score ranges were defined as follows:

- top range (7 – 9)
- middle range (4 – 6)
- bottom range (1 – 3)

Should the panel fail to reach agreement using the first option, the second option was to allocate the items in two sets and allocate items with a median score of 7 or above and an agreement level greater than 70% to the first set, and all other items to the second set.

Once data collection was completed, the ARRIVE working group met via videoconference to review the results and discuss the allocation of items into sets. As only 10 of 22 items reached the predefined agreement consensus of 70% (see supplementary information S3 – Delphi results), the second option was used to allocate items into two sets.

### 2 Results

#### 2.1 Composition of the Delphi expert panel

One hundred experts were invited to participate in the Delphi exercise, 73 accepted the invitation and 71 participated in the final round (see Figure 1). Ten data entries, which did not follow data dispersal rules were excluded, 61 data entries were therefore included in the final score analysis.

**Figure 1.**
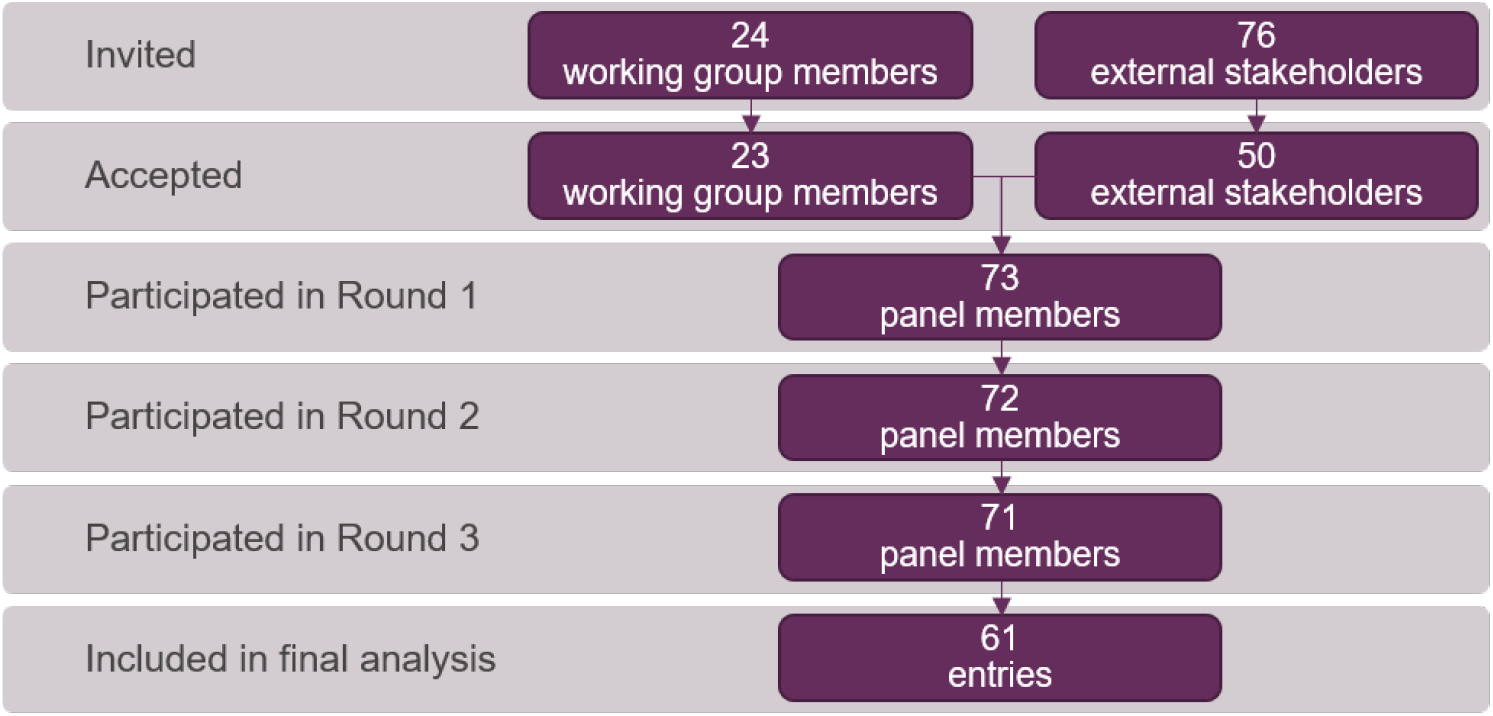
Delphi panel included at each stage of the Delphi exercise.

Demographics of the panel are presented in Table 2.

#### 2.2 Suggestions for new items and feedback on existing items

18 panel members suggested a total of 31 new items (see supplementary information S3 – Delphi data). No new item suggestion met the 10% threshold for inclusion in the revised guidelines.

Feedback on the wording of existing items was considered by the working group in the drafting of the revised items and the drafting of the accompanying E&E document.

Feedback from the Delphi panel indicated that the item on number analysed was misunderstood and confused with the item on sample size. For clarity, the item on number analysed was incorporated to the item on inclusion and exclusion criteria in further iterations of the guidelines. This reduced the number of items to 21.

#### 2.3 Scores for each Delphi round

The scores assigned to each item in rounds 1, 2 and 3 are shown in Figure 2.

**Figure 2.**
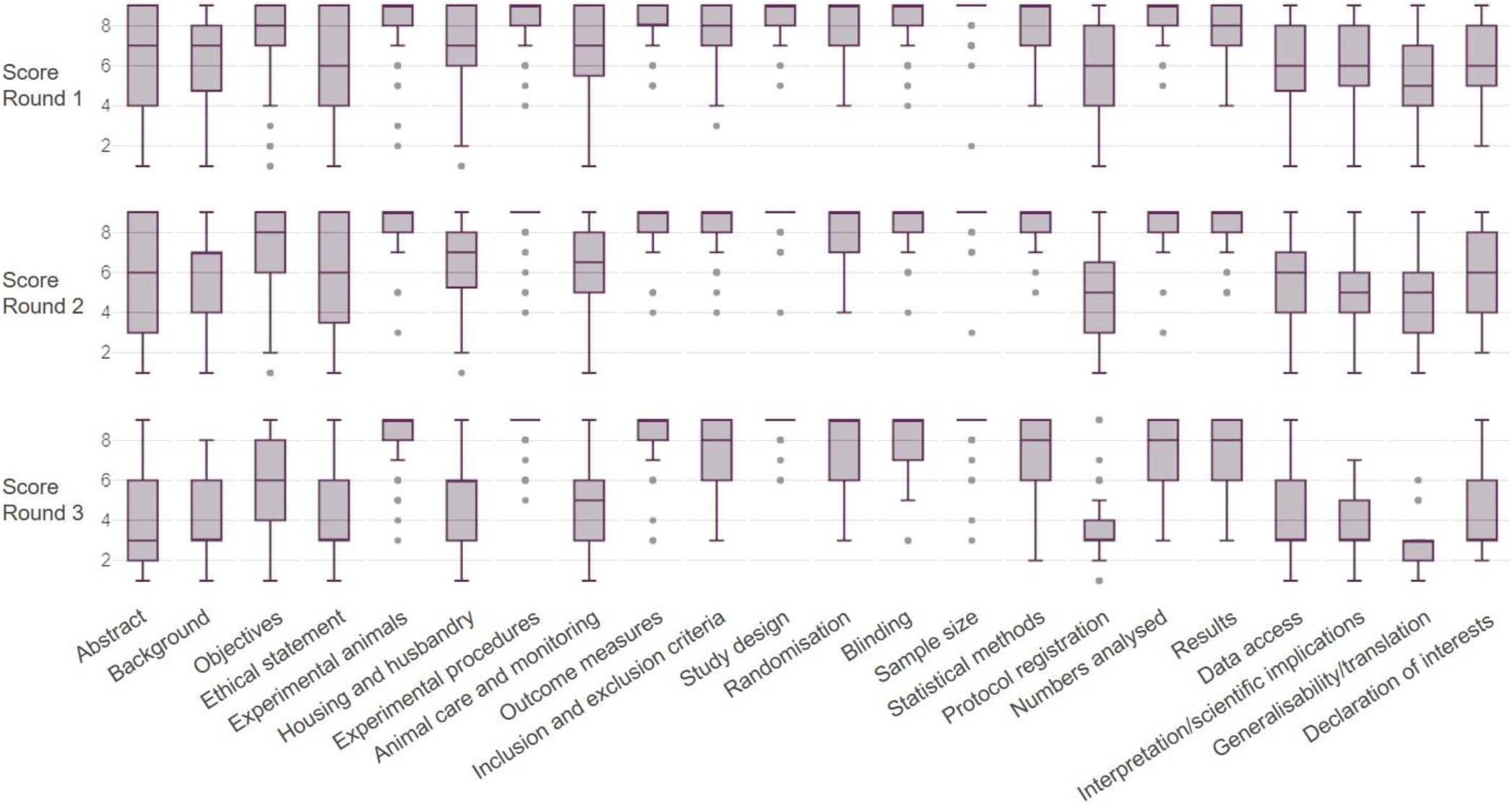
Item scores for each of the three Delphi rounds. Box and whisker plots of the panel members’ scores for the 22 items. Round 1: n=71−73, round 2: n=70−71, round 3: n=61, the exact sample size for each item in each round is provided in supplementary information S3 – Delphi data. Data plotted as median, interquartile range, minimum, maximum and outliers using https://www.displayr.com/. Raw data available at https://osf.io/8xjdr/.

#### 2.4 Allocation of items to sets

The allocation of items into sets is presented in Table 1. Eight items were shortlisted based on the a priori criterion (score in the top range and over 70% agreement within the panel). Three further items scoring in the top range were added to the shortlist following discussion within the working group.

**Table 1.** Allocation of the 22 items into two sets. Scores are displayed as median and interquartile range(IQR), n=61.

| Set | Item | Score | % scores in the top range | Reasoning |
| --- | --- | --- | --- | --- |
| Set 1 | Study design | 9 (9-9) | 95 | All items met pre-defined threshold (70%) for Set 1. |
|  | Sample size | 9 (9-9) | 92 |  |
|  | Experimental procedures | 9 (9-9) | 87 |  |
|  | Outcome measures | 9 (8-9) | 85 |  |
|  | Experimental animals | 9 (8-9) | 84 |  |
|  | Blinding | 9 (7-9) | 77 |  |
|  | Randomisation | 9 (6-9) | 70 |  |
|  | Statistical methods | 8 (6-9) | 72 |  |
|  | Numbers analysed | 8 (6-9) | 69 | Median score in the top range. |
|  | Inclusion and exclusion criteria | 8 (6-9) | 66 |  |
|  | Results | 8 (6-9) | 59 |  |
| Set 2 | Objectives | 6 (4-8) | 33 | Median score outside the top range. |
|  | Housing and husbandry | 6 (3-6) | 21 |  |
|  | Animal care and monitoring | 5 (3-6) | 10 |  |
|  | Ethical statement | 3 (3-6) | 20 |  |
|  | Abstract | 3 (2-6) | 16 |  |
|  | Data Access | 3 (3-6) | 11 |  |
|  | Background | 3 (3-6) | 8 |  |
|  | Protocol registration | 3 (3-4) | 7 |  |
|  | Interpretation/Scientific implications | 3 (3-5) | 2 |  |
|  | Declaration of interests | 3 (3-6) | 2 |  |
|  | Generalisability/Translation | 3 (2-3) | 0 |  |

**Table 2.**
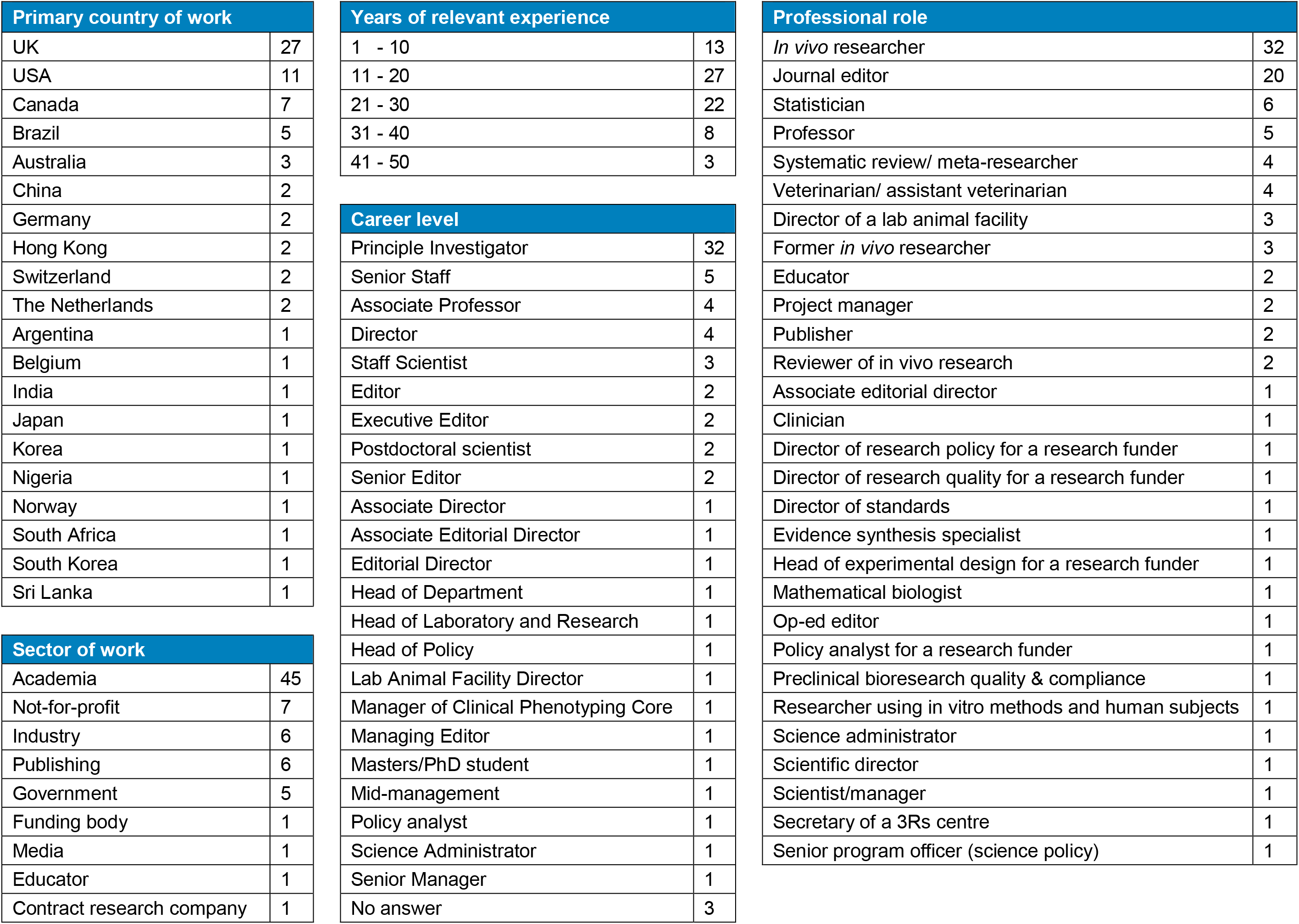
Demographics of Delphi respondents (n = 73). Note that the total number of professional roles exceeds the number of panel members as they could select more than one role.

Note that 11 items were allocated to set 1 but the combination of inclusion and exclusion criteria and numbers analysed in subsequent iterations of the guidelines reduced that number to 10 shortlisted items.

